# Structural Hijacking of FcRn by Human Astrovirus Spikes Reveals Conserved Epitopes for Broad-Spectrum Antivirals

**DOI:** 10.1101/2025.09.13.676064

**Authors:** Sashank Agrawal, Monika Jain, Daniella Marinelli, Bryan Briney, Ian A. Wilson

**Affiliations:** Department of Integrative Structural and Computational Biology, The Scripps Research Institute; La Jolla, CA, 92037 USA; Department of Immunology and Microbiology, The Scripps Research Institute, La Jolla, CA, 92037 USA; The Skaggs Institute for Chemical Biology, The Scripps Research Institute; La Jolla, CA, 92037 USA

## Abstract

Human astroviruses (HAstVs) are a leading cause of pediatric gastroenteritis and emerging systemic infections, yet no targeted therapies exist. A critical barrier to intervention has been the lack of molecular insights into viral entry, particularly the interaction between the HAstV capsid spike and its receptor, the neonatal Fc receptor (FcRn). Here, we report high-resolution crystal structures of the HAstV spike from classical serotypes 2 and 6 in complex with human FcRn, defining the conserved receptor-binding interface at atomic resolution. These structures reveal serotype-specific variations dictating receptor affinity and that reported neutralizing antibodies inhibit infection primarily by sterically blocking the receptor-binding site. Mapping conserved epitopes across classical HAstV serotypes provides a blueprint for design of broad-spectrum antivirals that disrupt viral entry. Notably, our structural data rationalize the potential repurposing of clinical FcRn inhibitors, such as Nipocalimab, to block HAstV infection, bridging critical gaps in astrovirus biology and antiviral development.

## Introduction

Human astroviruses (HAstVs) are the leading cause of viral gastroenteritis cases worldwide, particularly affecting young children, the elderly, and immunocompromised individuals.^1–4^ In addition to gastrointestinal illness, HAstVs have also been implicated in more severe extraintestinal infections, including meningitis and encephalitis, highlighting the virus’s potential for significant morbidity beyond the gut.^5–8^ Among the eight classical serotypes (HAstV-1 to HAstV-8), HAstV-1 is the most prevalent.^9–12^ The mature viral capsid, formed by three structural proteins derived from proteolytic cleavage of the ORF2-encoded precursor, features around 30 homodimeric spikes composed of VP27 subunits compared to the immature virion with 90 VP27 spikes.^13^ These spikes mediate host cell attachment and are the primary target of neutralizing antibodies (nAbs).^14^ Neutralizing antibodies against classical astroviruses are prevalent among adults, with most individuals exhibiting immunity to at least one serotype.^9,15^ This widespread seropositivity suggests that most people are exposed to astroviruses at some point in their life, typically by adulthood. Despite their high seroprevalence and clinical significance, how HAstV gains entry in host cells remained unresolved until the recent identification of the neonatal Fc receptor (FcRn) as the functional receptor for classical strains.^16,17^

FcRn, encoded by the FCGRT gene, is a heterodimeric receptor with structural homology to MHC class I molecules.^18^ Comprised of an α-chain (α1, α2, α3 domains) and β2-microglobulin (β2m), FcRn is best known for regulating IgG homeostasis. FcRn binds IgG in acidic endosomes (pH ≤ 6.5) and releases it at neutral pH (7.4), thereby extending antibody half-life and facilitating transcytosis across epithelial barriers.^19^ Recent studies revealed that classical HAstVs directly engage FcRn via their VP27 spike protein to initiate infection.^16,17^ While this discovery clarified HAstV tropism, the molecular architecture of the spike-FcRn interface remains undefined, hindering the development of cross-reactive vaccines or therapies targeting conserved epitopes.

Structural characterization of viral-receptor complexes is critical for deciphering entry mechanisms and guiding therapeutic development. High-resolution structures of the HAstV spike alone revealed a globular β-barrel core and hypervariable surface loops. However, how the virus targets FcRn is not known due to the lack of a spike-FcRn complex structure.^20^ While several neutralizing antibodies targeting HAstVs have been identified, their mechanisms of action are also not fully elucidated. It is unclear whether neutralizing antibodies block infection by directly competing with FcRn binding, by allosterically destabilizing the spike-receptor interaction, or by other mechanisms. This knowledge gap complicates vaccine design, as current strategies often prioritize immunodominant but variable loops, which drive strain-specific immunity, over conserved, functionally critical epitopes. Resolving the spike-FcRn interface is essential to shift the focus toward conserved epitopes that confer broad protection across diverse HAstV strains, enabling development of more universal therapies and vaccines. Furthermore, recent work has demonstrated that anti-FcRn antibodies, such as Nipocalimab, can block HAstV infection in cell lines and in *ex vivo* human enteroid models.^17^ Thus, deciphering the structural basis for this inhibition could provide insights into therapeutic interventions.

Here, we present high-resolution crystal structures of the HAstV spike from two different serotypes, 2 and 6, in complex with human FcRn. These structures define the atomic basis for receptor recognition, revealing a conserved receptor binding interface centered on the spike’s β-barrel core. This interface explains serotype-specific differences in receptor affinity and pinpoints residues critical for FcRn engagement. These structural insights demonstrate that most neutralizing antibodies to date act through steric competition rather than allosteric modulation, directly occluding the FcRn-binding site. Furthermore, by mapping conserved epitopes across classical serotypes, this work provides a blueprint for designing biologics or small-molecule inhibitors that broadly disrupt HAstV entry by blocking virus-receptor interaction. Importantly, our structural data support the potential repurposing of clinical FcRn inhibitors as a therapeutic strategy for HAstV. By bridging gaps in astrovirus entry mechanisms and FcRn biology, this work advances strategies to combat HAstV and related enteric pathogens through structure-guided interventions.

## Results

### Classical HAstV Spikes Bind FcRn Across Physiological pH Ranges

To characterize the interaction between classical HAstV spike proteins and the FcRn receptor, we expressed and purified the untagged recombinant human FcRn extracellular domain and Avi-tagged spike proteins from HAstV serotypes 1, 2, 6, and 8 in human 293 cell lines (see methods). Using biolayer interferometry (BLI), we immobilized biotinylated spikes on streptavidin-coated biosensors and measured their binding kinetics against the FcRn protein. These experiments confirmed direct, high-affinity interactions between HAstV spikes and FcRn, with equilibrium dissociation constants, Kd, in the nanomolar range across physiological pH conditions (Figure 1A).

**Figure 1.**
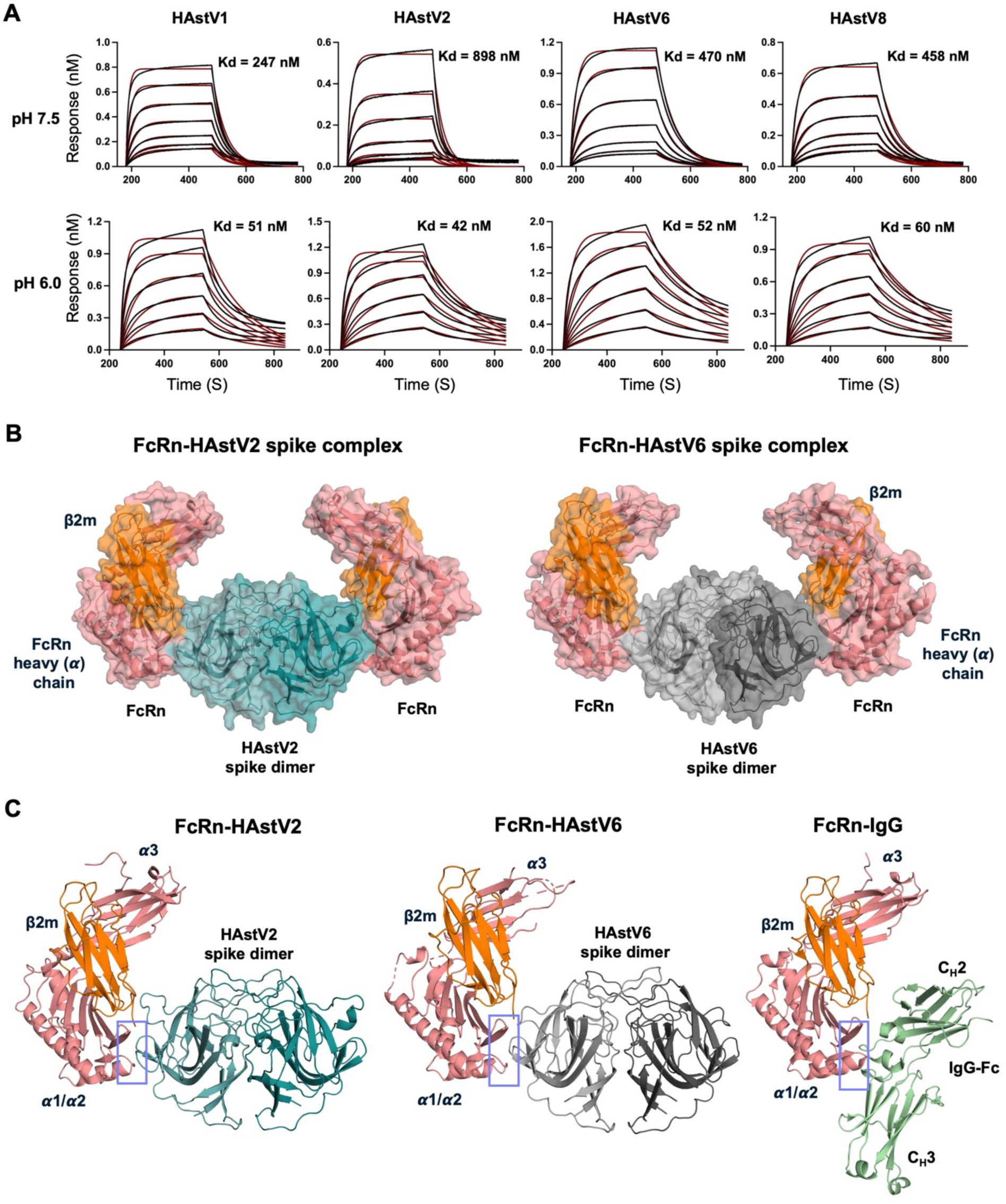
FcRn binds classical HAstV spikes with 2:2 stoichiometry. **(A)** Binding affinity of HAstV1, HAstV2, HAstV6, and HAstV8 spikes to FcRn measured by biolayer interferometry (BLI) at pH 6.0 and 7.5. Red curves show fitted data and black lines represent experimental data. **(B)** Crystal structure of FcRn in complex with HAstV2 and HAstV6 spikes, illustrating 2:2 stoichiometry (one FcRn per spike protomer). FcRn heavy chain (salmon), β2m (orange), HAstV2 (cyan), and HAstV6 (grey) are shown. **(C)** Structural comparison of FcRn bound to HAstV2 (left), HAstV6 (middle), and IgG-Fc (right, PDB: 6WOL), demonstrating that HAstV spikes and IgG-Fc bind the same region on FcRn α1/α2 domains. Only one FcRn is shown per spike dimer for clarity. The interface is marked with a blue box.

Since FcRn binds IgG exclusively at acidic pH (≤6.5) but not at neutral pH (7.4), we investigated whether HAstV spike-FcRn interactions display similar pH dependence.^21^ Notably, HAstV spikes exhibited significant binding at both neutral (pH 7.5) and acidic (pH 6.0) pH, contrasting sharply with FcRn’s pH-dependent IgG binding. At neutral pH, spikes demonstrated high nM binding with Kd values of 247 nM (serotype 1), 898 nM (serotype 2), 470 nM (serotype 6), and 458 nM (serotype 8), which increased substantially at lower pH (Kd values: 51 nM [serotype 1], 42 nM [serotype 2], 52 nM [serotype 6], and 60 nM [serotype 8]). This enhanced affinity at low pH was driven primarily by slower dissociation rates (Figure 1A and table S1). The retention of HAstV spike-FcRn binding across physiological pH suggests that HAstV leverages FcRn for initial cell surface attachment and viral entry throughout the variable intraluminal pH conditions of the human intestine.^22–24^ Stronger binding at low pH may further facilitate endosomal retention and viral uncoating. These findings establish FcRn as a functional HAstV receptor and reveal a binding mechanism that diverges from FcRn’s classical IgG-binding mode, which is strictly pH-dependent.

### FcRn Recognition by HAstV Spikes Shows a Binding Interface Shared with IgG Fc, but with a Divergent Mechanism

To elucidate the structural basis of HAstV spike interaction with FcRn, we determined crystal structures of HAstV2 and HAstV6 spikes in complex with FcRn. The HAstV2-FcRn complex diffracted to 3.07 Å resolution, while the HAstV6-FcRn complex to 2.97 Å resolution. In the HAstV2-FcRn structure, the asymmetric unit contains FcRn and spike in a 2:2 stoichiometry, forming a biological assembly in which two FcRn heterodimers (α-chain and β2-microglobulin) symmetrically engage a spike dimer. Each FcRn α-chain binds one protomer of the spike dimer (Figure 1B). In the HAstV6-FcRn structure, the asymmetric unit contains a single copy HAstV spike protomer and one FcRn heterodimer where the 2:2 biological assembly is crystallographic, with FcRn and spike arranged in the same way as the HAstV2-spike complex (Figure 1B). In both structures, FcRn engages the HAstV spike primarily through its α1/α2 domains, which are the same regions responsible for IgG Fc binding, with negligible direct contributions from β2m (Figure 1C).^25^

Comparative structural analysis of the FcRn–spike and FcRn–IgG complexes demonstrates that they share overlapping interface residues (Figure 2A). Buried surface area analysis further supports this observation, showing that several FcRn residues are in common at the interface of both complexes (Figure 2B). Notably, key residues involved in IgG binding also mediate interactions with the HAstV spikes, indicating a shared binding interface. In particular, Glu115, Glu116, and the 130DWPE133 motif within the α1/α2 domain of FcRn are critical for both FcRn–IgG and FcRn–spike interactions (Figures 2B, 2C, and 3). The buried surface area (BSA) for the HAstV2-FcRn complex is 508 Å² for FcRn and 523 Å² for the spike, whereas the HAstV6-FcRn complex has a larger BSA of 607 Å² for FcRn and 625 Å² for the spike, reflecting enhanced structural engagement that correlates with higher binding affinity. For comparison, the BSA in the human FcRn–IgG complex (PDB: 6WOL) is 732 Å² for FcRn and 665 Å² for IgG.^25,26^

**Figure 2.**
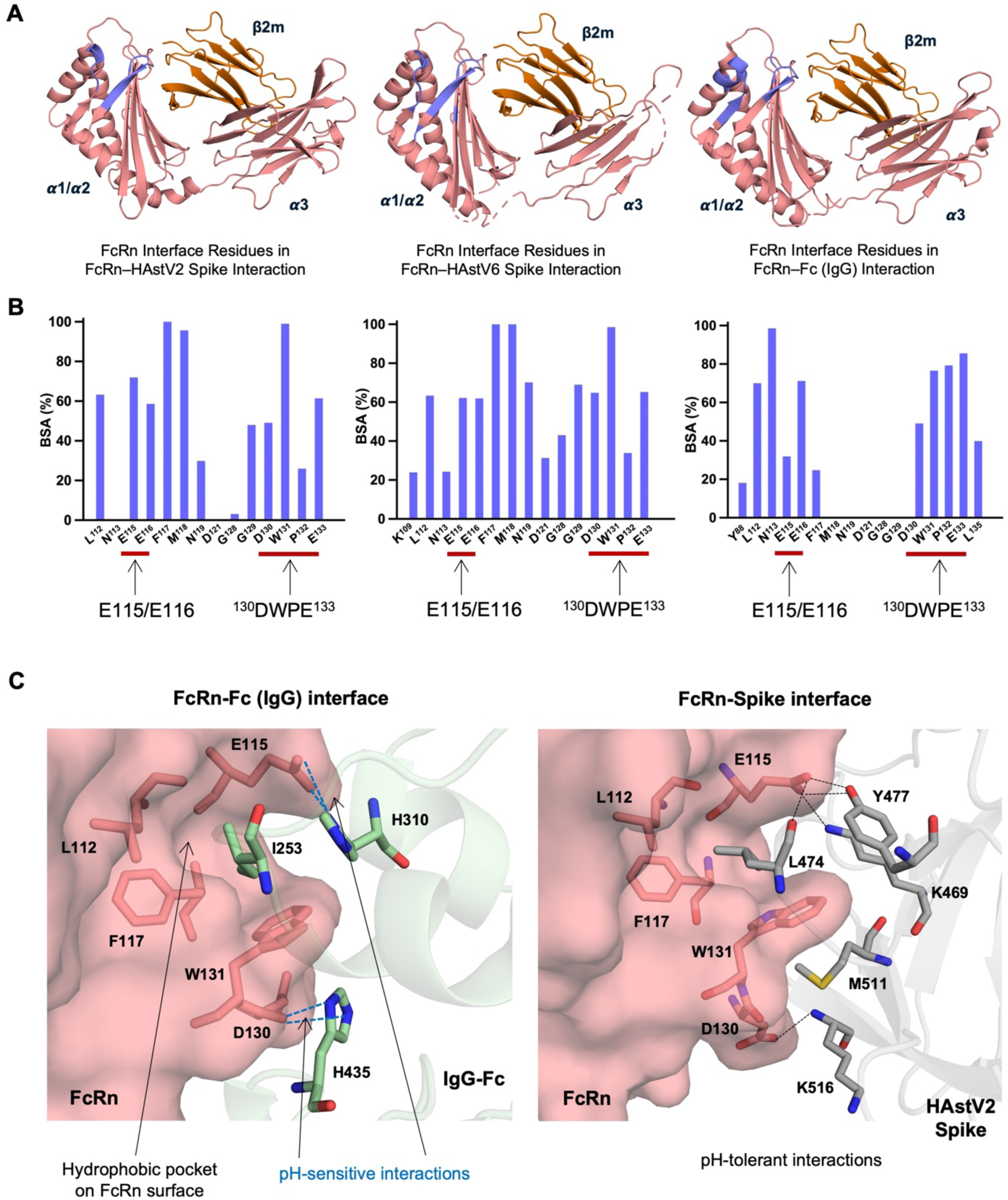
HAstV spikes and IgG-Fc share a common binding site on FcRn. **(A)** Cartoon representation of FcRn in complex with HAstV2, HAstV6, and IgG-Fc, with interface residues colored blue to highlight the shared interaction interface. FcRn heavy chain (salmon) and β2m (orange) are shown. **(B)** Buried surface area (BSA) of FcRn residues involved in binding HAstV2, HAstV6, and IgG-Fc, shown as a percentage of total buried area per residue. Shared interaction sites on FcRn are marked with red lines. **(C)** Left: The FcRn-IgG interface (PDB: 6WOL) with FcRn shown as a salmon surface and IgG as a pale green cartoon. Key interacting residues are shown as sticks. IgG Ile253 anchors to FcRn’s hydrophobic pocket; pH-sensitive interactions between IgG His310/His435 and FcRn E115/D130 are indicated by dotted blue dashed lines. Right: FcRn surface (salmon) and Spike cartoon (gray) with interacting residues (sticks) that mediate binding. Here, HAVst2 Leu474 anchors to FcRn’s hydrophobic pocket. pH-tolerant interactions corresponding to the pH-sensitive FcRn-IgG interactions are indicated by black dashed lines.

While both IgG and HAstV spike engage overlapping regions on FcRn α1/α2 domains, their binding mechanisms diverge sharply. In contrast, albumin binds FcRn at a completely distinct, non-overlapping site.^25^ IgG relies on pH-dependent interactions, inserting its Ile253 into a hydrophobic pocket on FcRn and positioning its His310/His435 residues to form pH-sensitive ionic bonds with FcRn acidic residues, Glu115, Glu130 (Figure 2C).^18^ In contrast, the HAstV spikes bind FcRn in a pH-tolerant manner, mediated by a bipartite strategy combining hydrophobic anchoring and polar interactions (Figure 2C, and 3).

Although HAstV2 and HAstV6 are antigenically diverse with only 46% sequence identity at protein level, they engage FcRn in a similar fashion. Both crystal structures reveal a hydrophobic pocket in the HAstV spike that accommodates FcRn Trp131, in a critical interaction stabilized by a network of non-polar residues (Figure 3B). Key HAstV spike residues line this hydrophobic pocket, thereby engaging FcRn Trp131 through hydrophobic contacts while simultaneously forming polar interactions around the periphery (Figure 3C).

**Figure 3.**
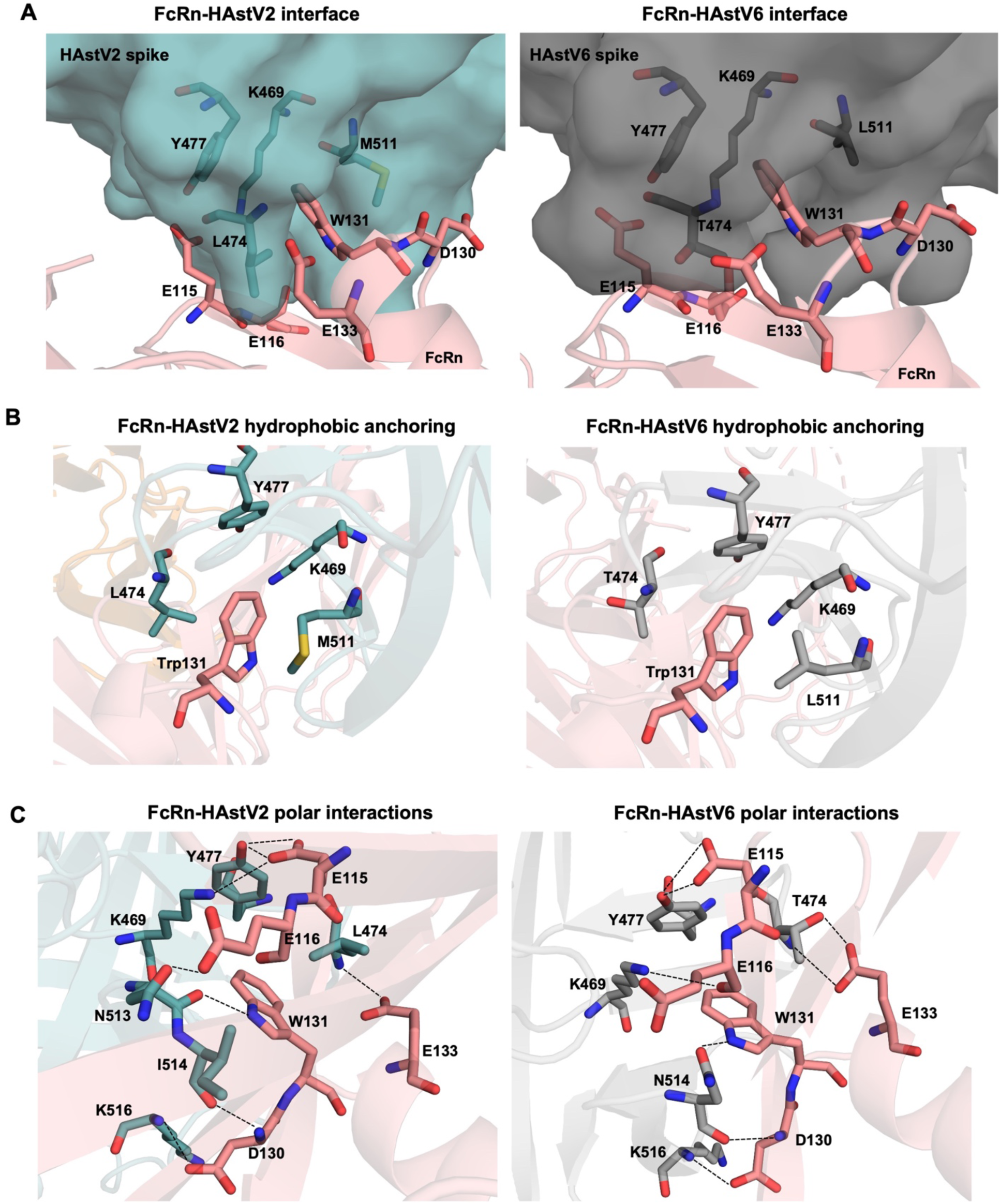
Structural basis of FcRn recognition by HAstV spikes. **(A)** Receptor-binding interface architecture in FcRn-HAstV2 and FcRn-HAstV6 spike complexes. Hydrophobic anchoring of FcRn W131 within conserved spike pockets and surrounding polar residues. Spike is shown as a surface, FcRn as a cartoon, and key residues as sticks. **(B)** Hydrophobic stabilization. Key residues (HAstV2: K469, L474, Y477, M511; HAstV6: K469, T474, Y477, L511) form van der Waals contacts with W131 in HAstV2 (left) and HAstV6 (right). **(C)** Polar interaction networks. Polar residues form salt bridges and hydrogen bonds to FcRn E115, E116, D130, and E133, reinforcing HAstV-FcRn complex interactions.

In the HAstV2–FcRn complex, the hydrophobic pocket is formed by the side chains of spike Lys469, Leu474, Tyr477, and Met511 (Figure 3B) (n.b. residue numbering is based on a sequence alignment of all classical HAstV spike proteins, anchored to the HAstV2 reference sequence due to its greater length as compared to other serotypes, as shown in Figure S1, ensuring consistent positional mapping across variants). Surrounding this pocket, several polar residues form hydrogen bonds and ionic interactions: Lys469 and Tyr477 interact with FcRn Glu115; Asn113 with FcRn Glu116 and Trp131; Lys516 and Ile514 with FcRn Asp130; and Leu474 with FcRn Glu133 (Figure 3C).

For the HAstV6-FcRn complex, the hydrophobic pocket is formed by spike Lys469, Thr474, Tyr477, and Leu511 (Figure 3B). Notably, Thr474 and Tyr477 play dual roles: their side chains contribute to the hydrophobic environment stabilizing Trp131 while also forming hydrogen bonds with FcRn Glu133 and Glu115, respectively. Additional polar interactions further reinforce the interaction, including Asn514 with FcRn’s Asp130 and Trp131, and Lys516 with FcRn Asp130 (Figure 3C). This hybrid binding strategy, which involves hydrophobic sequestration of Trp131 and a surrounding hydrogen-bond network, explains the high-affinity, pH-tolerant interaction, supporting binding at both acidic and neutral pH, and provides a structural rationale for serotype-specific variations in FcRn engagement.

### HAstV-Neutralizing Antibodies Sterically Block FcRn Binding Without Direct Epitope Overlap

While several neutralizing antibodies targeting classical human astroviruses (HAstVs) have been identified, their mechanisms of action remained poorly understood due to limited insight into virus-receptor interactions. To address this issue, we performed structural analyses of previously determined antibody-spike complexes in the context of the FcRn-spike binding.^14,27,28^ Notably, none of the known HAstV-neutralizing antibodies bind the precise FcRn interaction site on the HAstV spike protein. However, two antibodies, 3E8 and PL-2 (PDB: 7RK1 5KOV), partially overlap with regions of the FcRn binding site, particularly near the β5-β6 loop of the spike (Figure 4A and S1).^14,27^ For antibodies 3B4, 4B6, and 2D9, their epitopes do not directly overlap with FcRn’s binding footprint but instead engage adjacent regions of the HAstV spike (Figure 4A and S1).^14,28^ Structural modeling predicts that all five antibodies, including their single-chain variable fragments (scFvs), sterically clash with FcRn, preventing receptor engagement despite lacking direct epitope competition (Figure 4A and S2). In contrast, antibody 3H4 binds a distal epitope at the base of the HAstV spike, a region partially occluded by the adjacent capsid core domain in intact virions.^28,29^ To access this buried site, 3H4 likely induces conformational changes in the capsid or reorients the spike laterally, disrupting FcRn interactions through steric hindrance or allosteric effects. This mechanism distinguishes 3H4 from the other five antibodies, which block FcRn-spike binding directly through steric clash.

**Figure 4.**
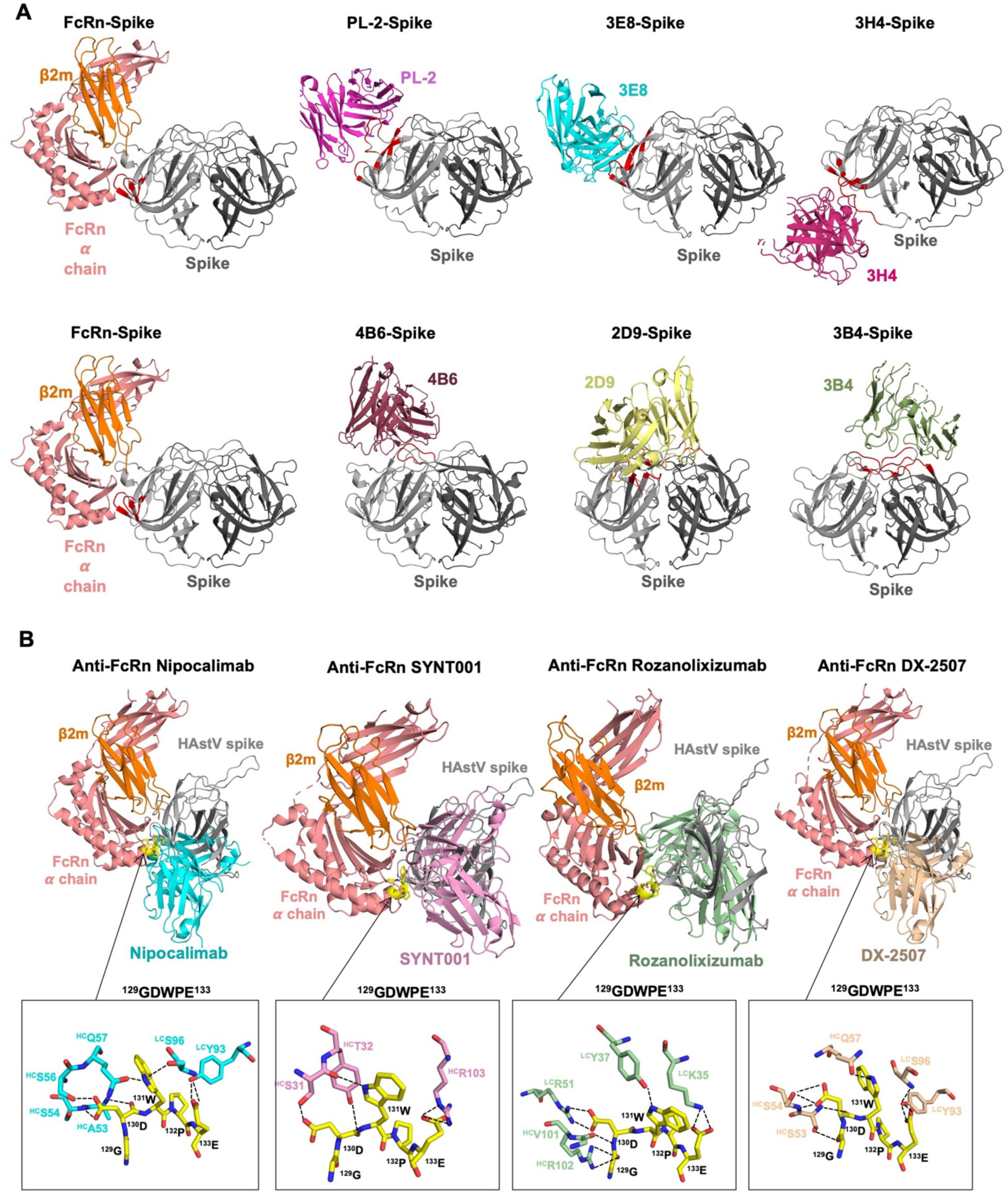
Structural basis for antibody neutralization and anti-FcRn antibody inhibition of HAstV infection. **(A)** HAstV neutralizing antibody interactions. Structural comparison of FcRn-spike complex with antibody-spike complexes: 3E8 (PDB: 7RK1), 2D9 (PDB: 7RK2), PL-2 (PDB: 5KOV), 3B4 (PDB: 9CBN), 4B6 (PDB: 9CN2), and 3H4 (PDB: 9CBN). All antibodies except 3H4 sterically block FcRn binding. 3H4 appears to neutralize through spike conformational changes rather than direct competition. Spike interface residues are colored red. **(B)** Mechanism for potential inhibition of HAstV infection by anti-FcRn antibodies. Structural alignment of FcRn-antibody complexes Nipocalimab (PDB: 9MI6), SYNT001 (PDB: 6NHA), DX-2507 (PDB: 5WHK), and Rozanolixizumab (PDB: 6FGB) with the FcRn-spike complex reveals direct epitope overlap. Residues ^129^GDWPE^133^ (yellow, inset) are targeted by all anti-FcRn antibodies shown.

### Structural Basis for Anti-FcRn Antibody Blockade of HAstV Infection

Recent studies demonstrated that the anti-FcRn monoclonal antibody Nipocalimab blocks classical HAstV infection in cell lines and ex vivo human enteroid models.^17^ To elucidate its mechanism of action, we performed structural alignment of the Nipocalimab-FcRn complex with our HAstV spike-FcRn complexes (Figure 4B). This analysis revealed overlapping binding interfaces at FcRn’s α1/α2 domains, indicating direct competition for receptor occupancy. This competitive inhibition is driven by Nipocalimab’s very high-affinity (Kd ≤57.8 pM at pH 7.4, and ≤31.7 pM at pH 6.0) and pH-independent interaction with FcRn,^26,30^ which forms an extensive interface (1017 Å²) surpassing that of both IgG-Fc (698 Å²) and HAstV spike complexes (647 Å² for HAstV6-FcRn, 548 Å² for HAstV2-FcRn). The antibody’s larger binding interface and strong affinity enables it to outcompete endogenous IgG and sterically block viral engagement of FcRn.

Comparative structural analysis of other high affinity anti-FcRn antibodies, SYNT001 (Kd 1.2 nM at pH 7.4), DX-2507 (2 nM), and rozanolixizumab (23 pM), reveals similar potential to disrupt HAstV infection.^31–34^ All of these anti-FcRn antibodies target FcRn residues that are critical for HAstV binding, particularly the ^129^GDWPE^133^ motif (Figure 4B). These FcRn inhibiting antibodies, either in clinical trials or approved for autoimmune disorders, bind FcRn with pH-independent, high-affinity interactions far exceeding the HAstV spike.^35–37^ Their larger binding interfaces and saturation of FcRn’s IgG-binding site likely prevent viral engagement with host cells. These findings highlight the therapeutic potential of repurposing FcRn-targeted antibodies to combat HAstV, a pathogen with no approved antivirals.

### Conserved Hydrophobic Pocket Mediates HAstV Spike-FcRn Binding Across Serotypes and Suggests Broad Therapeutic Targeting

The HAstV6 spike adopts a dimeric conformation, consistent with previously solved structures from other classical serotypes, HAstV1, HAstV2, and HAstV8. Structural alignment of HAstV6 with HAstV1 (Cα RMSD = 0.52 Å), HAstV2 (Cα RMSD = 0.64 Å), and HAstV8 (Cα RMSD = 0.47 Å) reveals a highly conserved fold across serotypes, with deviations localized primarily to flexible loop regions (Figure 5A, 5B).^14,20^ This structural homology underscores evolutionary conservation of the dimeric spike architecture, which likely serves as a functional prerequisite for receptor engagement and capsid assembly.

**Figure 5.**
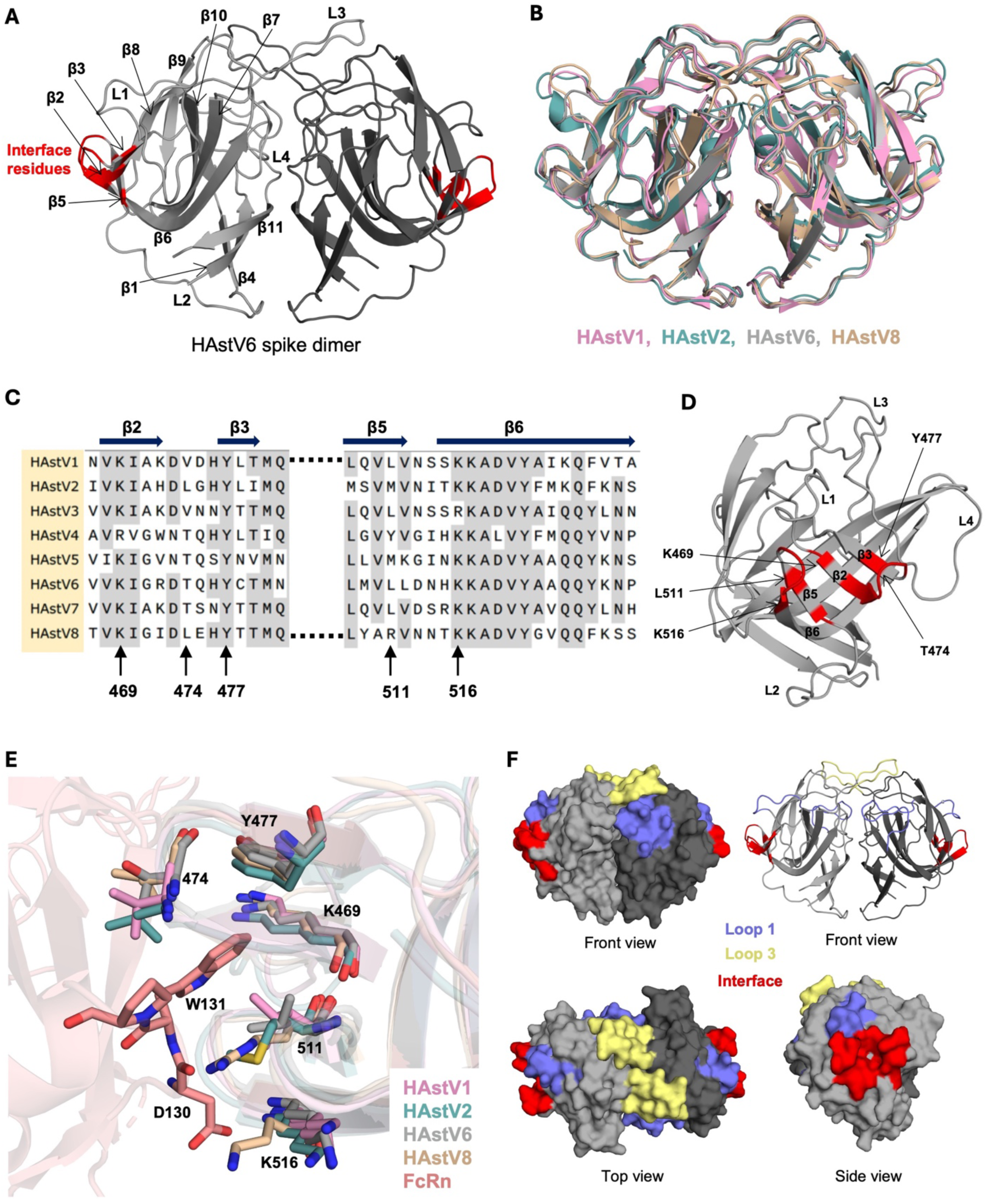
Structural conservation of HAstV spike dimers and FcRn-binding regions. **(A)** Dimeric architecture of the HAstV6 spike, with interface residues colored red and loop/beta sheet numbers shown on one protomer. **(B)** Alignment of HAstV6 spike dimer with HAstV1, HAstV2, and HAstV8, highlighting the conserved dimeric architecture. **(C)** Sequence alignment of classical HAstV spikes (only interface regions are shown). Key interacting residues are marked with arrows; beta sheet numbers are indicated above the sequence. **(D)** Side view of one HAstV spike protomer, with interface residues colored red and key residue locations indicated by arrows. **(E)** Structural alignment of HAstV2-FcRn and HAstV6-FcRn complex with HAstV1 (PDB:5EWO) and HAstV8 (PDB:7RK1) spikes, with key residues shown as sticks. **(F)** Front, top, and side views of the spike dimer, showing location and surface exposure of loop 1 (blue), loop 3 (yellow), and FcRn-binding residues (red).

The HAstV spike-FcRn interface is primarily stabilized by four hydrophobic residues on the spike that contact FcRn Trp131. Among these, Lys469 and Tyr477 are fully conserved across all eight classical HAstV serotypes, preserving a critical hydrophobic core as well as surface polarity (Figure 5C). Thr474, conserved in four serotypes (including HAstV6), is substituted by Val/Leu in others (including HAstV2) in a change that retains hydrophobicity but eliminates the hydrogen bonds made by the threonine OH group, as observed in HAstV6 (Figure 3C). Similarly, Leu511, conserved in four serotypes, is replaced by Met/Tyr/Arg in others; these substitutions maintain hydrophobic stacking with Trp131 through nonpolar side chains or atoms, as observed in the HAstV2-FcRn complex (Figure 3B, 5C). Strikingly, Lys516, which forms a salt bridges with FcRn Asp130, is also conserved across all classical HAstV spikes. Together, variations in these interfacial residues, coupled with differences in peripheral hydrogen bond networks, likely drive serotype-specific binding affinities.

Structural alignment of HAstV2 and HAstV6 spike-FcRn complexes with previously determined HAstV1 and HAstV8 spikes reveals that all four critical residues (Lys469, Lys474, Tyr477, Tyr 511), along with Lys516, are spatially conserved and positioned to contact key FcRn interface residues, including Trp131 (Figure 5E). This conservation, despite low sequence identity among serotypes, highlights a shared evolutionary strategy where HAstV spikes form a conserved hydrophobic pocket to engage FcRn Trp131 while leveraging a polar network in FcRn (Glu115, Glu116, Asp130, and Glu133) for binding specificity and orientation. The preservation of these critical residues and interactions across serotypes, even as peripheral residues diverge, provides the basis for FcRn engagement in cellular entry of HAstV.

## Discussion

This study provides a comprehensive structural and mechanistic framework for understanding HAstV interaction with FcRn, shedding light on key aspects of viral entry, antibody neutralization, and therapeutic intervention strategies. Our findings illustrate that HAstV engages FcRn at both neutral and more acidic pH, a feature distinct from the strict acidic pH dependence (<pH 6.5) observed for IgG binding. This finding immediately suggests a departure from the conventional FcRn-IgG interaction model and implies a distinct mechanism by which HAstV exploits this receptor. Supporting this notion, astrovirus primarily targets the epithelial cells of the upper small intestine,^38,39^ where the pH varies, ranging from approximately 6.0 in the upper section, to 7.1 in the midsection, and reaching 7.4 in the terminal ileum.^22–24^ This physiological pH gradient aligns with our finding that the HAstV spike protein binds more strongly to FcRn at pH 6.0, which corresponds to the intraluminal pH of upper small intestine.

The crystal structures of the HAstV2 and HAstV6 spike in complex with FcRn confirm a shared binding interface with IgG on FcRn α1/α2 domains but a divergent binding mechanism. This recognition mode is very different from a model proposed from molecular docking.^16,17^ HAstV employs a bipartite strategy, relying on hydrophobic anchoring and polar interactions to achieve binding across physiological pH. Unlike IgG, which binds FcRn primarily in acidic endosomal compartments, HAstV can potentially interact with FcRn across the range of intraluminal pH values found in the small intestine. This suggests that HAstV utilizes FcRn for initial attachment to cells, a crucial step in initiating infection, while its enhanced low-pH affinity could plausibly facilitate viral uncoating in endosomes, as supported by analogous findings in other viruses. For example, FcRn acts as both an attachment and uncoating receptor for several echoviruses, such as E18 and Echo 6, where it triggers uncoating at low endosomal pH.^40,41^ Similarly, in porcine reproductive and respiratory syndrome virus (PRRSV), FcRn promotes uncoating by interacting with the viral PRRSV N and M proteins at acidic conditions.^42,43^ Notably, dipeptidyl-peptidase IV (DPP4) has been identified as a potential cofactor in HAstV entry with no direct binding HAstV spike, although its role appears secondary to FcRn.^17^ The interplay between FcRn and DPP4 warrants further investigation to determine potential synergistic effects on viral entry.

The identification of key residues forming the conserved hydrophobic pocket, along with the network of polar interactions, provides a detailed structural explanation for its particular binding mode. Structural alignment revealed a conserved dimeric architecture, consistent across classical HAstV serotypes. Key residues Lys469 and Tyr477, within the hydrophobic pocket accommodating FcRn Trp131, are fully conserved, while variations in other interfacial residues and hydrogen bond networks likely mediate serotype-specific binding affinities and tropism. These finding highlights both conserved and variable features of the HAstV-FcRn interaction. The conserved hydrophobic core within the HAstV spike receptor-binding site, critical for FcRn recognition, presents a promising target for small-molecule antiviral development. This evolutionary constraint on the hydrophobic core, necessitated by its role in maintaining the spike-FcRn interface, likely limits viral escape via mutation, making it an attractive site for broad-spectrum inhibitors. Notably, the divergent HAstV strains (e.g., VA1, MLB1) exhibit structurally distinct spikes lacking classical receptor-binding motifs, suggesting alternative entry mechanisms that may bypass FcRn or utilize different receptors altogether.^44,45^ Further research is needed to determine the prevalence and mechanisms of these alternative entry pathways.

The HAstV spike protein exhibits pronounced sequence and structural diversity in its surface-exposed loop regions, which serve as primary targets for neutralizing antibodies.^27,28^ Hypervariable loops 1 (residues 440–460) and 3 (residues 560–570) undergo frequent amino-acid substitutions and conformational rearrangements across serotypes, driving immune evasion. In contrast, loop 4 stabilizes the dimeric interface, while loop 2 anchors the base of the spike near the capsid shell (Figure 5A, 5F). Notably, our findings on neutralizing antibodies provide critical insights into intervention strategies. While current antibodies, such as 4B6, 3E8, and 2D9, bind hypervariable loops, none directly engage the FcRn interaction site. For example, antibody 4B6 targets the tip of loop 3 in HAstV-2, engaging D564 and N565 through an interaction network that can be disrupted by escape mutations (e.g., D564E/N565D).^46^ Similarly, HAstV-8-specific antibodies 3E8 and 2D9 recognize overlapping loop 1 epitopes, where substitutions including Y464H/D597Y enable evasion.^46^ Serotype specificity arises from high sequence divergence in these loops (<40% identity between serotypes), restricting cross-reactivity. Structural flexibility further limits broad protection, as mutations (e.g., Ser463Pro in loop 1) disrupt antibody binding without compromising spike function.^47^ This observation underscores the importance of conserved structural elements as targets for next-generation therapies.

To address HAstV antigenic diversity, our structural data highlight conserved β-sheet motifs (β5-6, β2-3, β8) adjacent to the FcRn-binding site as promising targets for cross-serotype immunity. These regions exhibit structural stability and conservation across serotypes while maintaining surface exposure for antibody engagement (Figure 5C, 5D, 5F). Structural analyses suggest that antibodies targeting β5-6 or β2-3 could sterically block FcRn binding through spatial interference, even in single-chain variable fragment (scFv) formats, and thus may have greater potential to fully block FcRn binding compared to antibodies that bind other regions. This strategy exploits structural constraints imposed by receptor-binding requirements, as these regions are critical for maintaining the FcRn interaction interface. Redirecting immune responses toward these conserved, functionally essential regions could minimize viral escape and enhance cross-serotype efficacy, offering a blueprint for universal HAstV vaccines or biologics.

Consistent with recent studies demonstrating that the anti-FcRn antibody Nipocalimab blocks HAstV infection, our structural analysis supports a mechanism of direct competition for receptor occupancy. The structural alignment showing overlapping binding interfaces between Nipocalimab and the HAstV spike on FcRn, combined with the high-affinity and pH-independent binding of Nipocalimab, provides a strong rationale for its efficacy. The fact that other anti-FcRn antibodies (SYNT001, DX-2507, rozanolixizumab) share this potential further strengthens the case for repurposing these clinically approved drugs to combat HAstV infections, which is a particularly attractive approach given their established safety profiles and potential for rapid clinical translation.

In conclusion, our study provides a comprehensive understanding of the HAstV-FcRn interaction, defining a largely pH-tolerant binding mechanism distinct from IgG-FcRn, identifying conserved structural motifs as vulnerable sites for cross-serotype immunity, and revealing how neutralizing antibodies exploit steric competition rather than direct receptor-site engagement. The conserved hydrophobic core of the HAstV spike, critical for FcRn recognition, emerges as a prime target for small-molecule inhibitors that could disrupt viral entry with minimal risk of immune evasion. While our work focused here on classical HAstV serotypes, the entry pathways of divergent strains (e.g., VA1, MLB1) remain unresolved. Future efforts should prioritize evaluating anti-FcRn antibodies (e.g., rozanolixizumab) in HAstV infection models, resolving entry pathways in non-classical strains, and characterizing FcRn’s interplay with cofactors such as DPP4. By bridging high-resolution structural insights with translational applications, this work advances a roadmap for combating HAstV infections and related enteric pathogens through precision intervention strategies that exploit conserved viral vulnerabilities.

## Material and Methods

### Expression and purification of recombinant HAstV capsid spike proteins

Codon-optimized sequences encoding the HAstV capsid spike domains (HAstV-1 [AAC34717.1, residues 429–645], HAstV-2 [Q82446.1, residues 428–647], HAstV-6 [AZB52207.1, residues 429–643], HAstV-8 [Q9IFX1, residues 424–645]) containing C-terminal AviTag and His10X affinity tags were synthesized in pcDNA3.1 vector. For transfection, 800 µg of HAstV spike plasmid and 250 µg of BirA plasmid were diluted in 100 mL OptiMEM serum-free media (Life Technologies, 31985070), sterile-filtered (0.22 µm SteriFlip, Millipore Sigma SCGP00525), and complexed with 800 µL FectoPRO transfection reagent (Polyplus, 116-040) for 15 min. The mixture was added to Expi293F cells (Thermo Scientific, A14527) at 3 × 10⁶ cells/mL and cultured for 5 days (37°C, 8% CO₂, 225 RPM).

At 24 hr post-transfection, cultures were supplemented with 300 mM valproic acid (Sigma, P4543) and 45% glucose (Sigma, G8769). Cells were pelleted by centrifugation at 3,500 RPM on day 5, and supernatants were filtered (0.22 µm, Thermo Scientific 09-741-06). Filtered supernatants were incubated overnight at 4°C with Pierce High-Capacity Ni-IMAC resin (Thermo Scientific; 500 µL resin per 1 L supernatant). Proteins were purified using a gravity flow column (Takara Bio, 635513), buffer-exchanged into 1× PBS (Corning, 21-030-CM) using Amicon Ultra 15 mL centrifugal filters (30 kDa MWCO; Millipore UFC9030), and further purified by size-exclusion chromatography (Superdex 200 column, GE Healthcare) on an AKTA Express system (GE Healthcare).

### Expression and purification of untagged recombinant human FcRn ectodomain

Human FcRn α-chain (residues 24-297; Uniprot P55899) and β2-microglobulin (residues 21-119; Uniprot P61769) were cloned into separate phCMV3 vectors. Plasmid DNA encoding FcRn heavy (α) and light (β2-m) chains (1:1 ratio) were co-transfected into Expi293F cells using ExpiFectamine 293 Transfection Kit in OptiMEM media. At 24 hr post-transfection, cultures were supplemented with 300 mM valproic acid and 45% glucose. Cultures were harvested at day 6 by centrifugation (3,000 × g, 20 min) and sterile filtered through a 0.22 µm filter. FcRn was purified from the supernatant using a human IgG column and further purified by size-exclusion chromatography on a Superdex 200 Increase 10/300 GL column (Cytiva 28-9909-44), pre-equilibrated with TBS at pH 7.4. Fractions corresponding to FcRn were collected and analyzed by SDS-PAGE to confirm purity.

### Biolayer interferometry (BLI) binding assays

Direct binding interactions between HAstV spike proteins (serotypes 1, 2, 6, 8) and human FcRn were analyzed using an Octet Red instrument (Sartorius). Biotinylated HAstV spike proteins (5 μg/mL in 1× kinetics buffer: PBS pH 7.4, 0.01% BSA, 0.002% Tween 20) were immobilized onto streptavidin (SA) biosensors. FcRn was tested at 2-fold serial dilutions (1,000 to 31.25 nM) at 23°C. Kinetic parameters were determined using a 1:1 binding model in Octet Analysis Studio software (Sartorius) with reference subtraction for baseline drift correction. The resulting association (k_a_), dissociation (k_dis_), and equilibrium dissociation constants (K_d_) are summarized in Table S1.

### Crystallization of FcRn-Spike complexes

FcRn and HAstV spike proteins were mixed at a 1:1 molar ratio in TBS buffer (pH 7.5), concentrated to 10 mg/mL, and incubated for 1 h at room temperature. Crystallization screening was performed using our CrystalMation high-throughput robotic system (Rigaku) with JCSG 1–4 and Top96 Cryo screens at 20°C. HAstV2-FcRn crystals grew in 25% 1,2-propanediol, 10% glycerol, 5% PEG-3000, 0.1 M phosphate-citrate (final pH 4.9). HAstV6-FcRn crystals were formed in 30% 1,2-propanediol, 20% PEG-400, 0.1 M HEPES (final pH 7.4). Crystals were cryoprotected by soaking in reservoir solution supplemented with 15% ethylene glycol and flash-cooled in liquid nitrogen for data collection.

### Structure determination and refinement

Diffraction data were collected at beamline 17-ID-2 (FMX) of the National Synchrotron Light Source II. Data indexing, integration, and scaling were performed using HKL2000.^48^ Structures were determined by molecular replacement in Phenix Phaser^49^ using initial models of FcRn (PDB 6WOL), HAstV2 spike (PDB 5KOU), and an AlphaFold 3-predicted model^50^ for the HAstV6 spike. Model building and iterative refinement were carried out using Coot and Phenix.refine, respectively.^51,52^ The quality of the final structures was assessed with MolProbity^53^ and further validated through the PDB validation server. Data collection and refinement statistics are summarized in Table S2.

## Acknowledgments

We thank H. Tien for assistance with the automated robotic crystal screening at The Scripps Research Institute. X-ray diffraction datasets were collected at the National Synchrotron Light Source II (NSLS II) beamline 17-ID-2. This research used resources of the National Synchrotron Light Source II, a US DOE Office of Science User Facility operated for the DOE Office of Science by Brookhaven National Laboratory under contract no. DE-SC0012704.

## Author contributions

S.A., B.B., and I.A.W. conceptualized the study. S.A. and D.M. prepared recombinant proteins. S.A. and M.J. crystallized samples, collected, and analyzed X-ray data. S.A. performed BLI experiments and prepared figures. S.A. and I.A.W. wrote the original manuscript. All authors reviewed and edited the manuscript. B.B. and I.A.W. acquired funding.

## PDB code availability

The x-ray coordinates and structure factors have been deposited in the Research Collaboratory for Structural Bioinformatics (RCSB) PDB under accession codes 9OC7 for the FcRn-HAstV2 spike complex, and 9OC6 for the FcRn-HAstV6 spike complex.

