## Supplementary material for "Structural Hijacking of FcRn by Human Astrovirus Spikes Reveals Conserved Epitopes for Broad-Spectrum Antivirals": Figure S1, Figure S2, Table S1, Table S2

for

**Affiliations**:

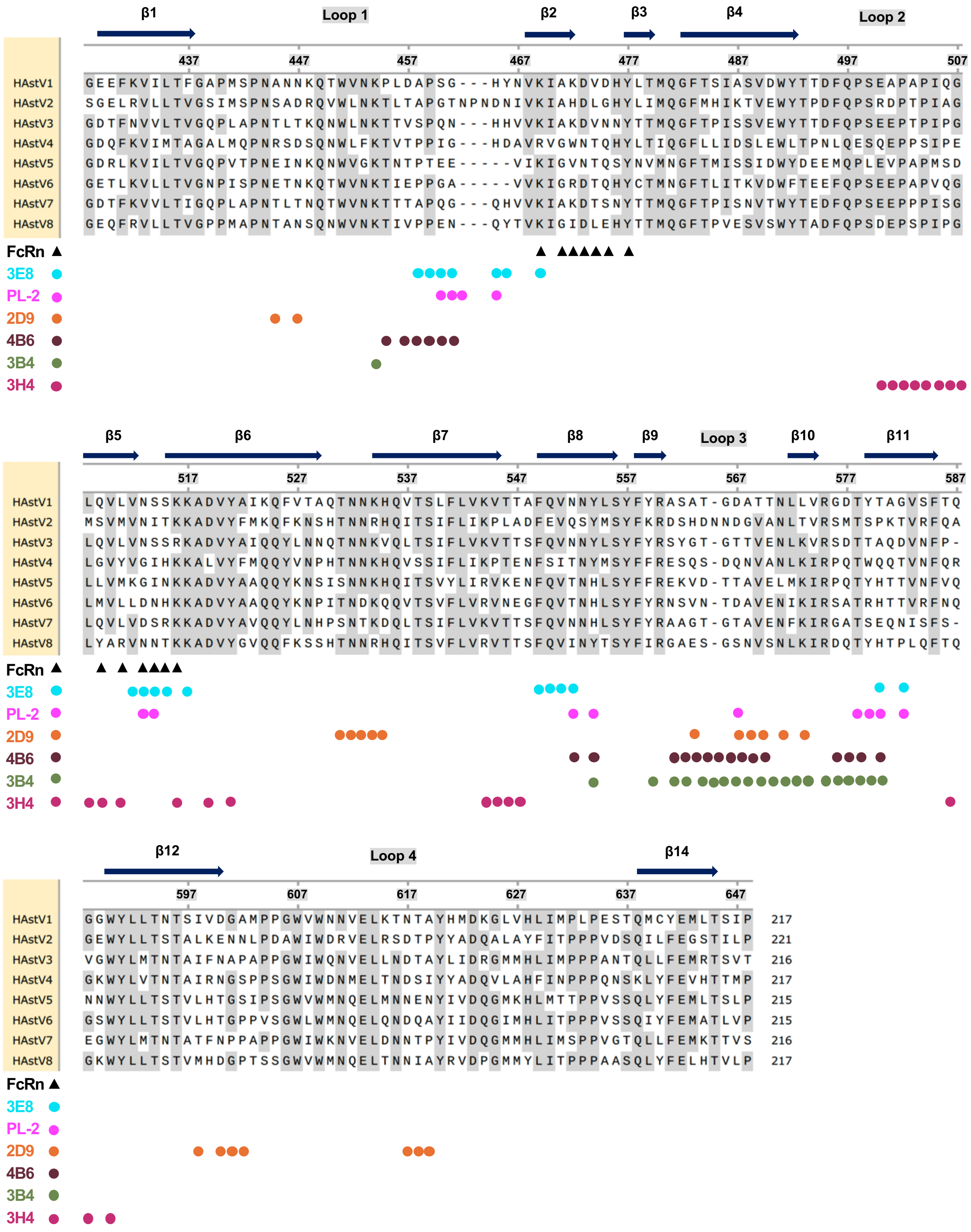
β1 Loop 1 β2 β3 β4 Loop 2
437 447 457 467 477 487 497 507
HAstV1 GEEFKVILTFGAPMSPNANNKQTWVNKPLDAPSG---HYNVKIAKDVDHYLTMQGFTSIASVDWYTTDFQPSEAPAPIQG
HAstV2 SGELRVLLTVGSIMSPNSADRQVWLNKTLTAPGTNPNDNIVKIAHDLGHYLIMQGFMHIKTVEWYTPDFQPSRDPTPIAG
HAstV3 GDTFNVVLTVGQPLAPNTLTKQNWLNKTTVSPQN---HHVVKIAKDVNNYTTMQGFTPISSVEWYTTDFQPSEEPTPIPG
HAstV4 GDQFKVIMTAGALMQPNRSDSQNWLFKTVTPPIG---HDAVRVGWNTQHYLTIQGFLLIDSLEWLTPNLQESQEPPSIPE
HAstV5 GDRLKVILTVGQPVTPNEINKQNWVGKTNTPTEE-----VIKIGVNTQSYNVMNGFTMISSIDWYDEEMQPLEVPAPMSD
HAstV6 GETLKVLLTVGNPISPNETNKQTWVNKTIEPPGA-----VVKIGRDTQHYCTMNGFTLITKVDWFTEEFQPSEEPAPVQG
HAstV7 GDTFKVVLTIGQPLAPNTLTNQTWVNKTTTAPQG---QHVVKIAKDTSNYTTMQGFTPISNVTWYTEDFQPSEEPPPISG
HAstV8 GEQFRVLLTVGPPMAPNTANSQNWVNKTIVPPEN---QYTVKIGIDLEHYTTMQGFTPVESVSWYTADFQPSDEPSPIPG
FcRn
3E8
PL-2
2D9
4B6
3B4
3H4
β5 β6 β7 β8 β9 Loop 3 β10 β11
517 527 537 547 557 567 577 587
HAstV1 LQVLVNSSKKADVYAIKQFVTAQTNNKHQVTSLFLVKVTTAFQVNNYLSYFYRASAT-GDATTNLLVRGDTYTAGVSFTQ
HAstV2 MSVMVNITKKADVYFMKQFKNSHTNNRHQITSIFLIKPLADFEVQSYMSYFKRDSHDNNDGVANLTVRSMTSPKTVRFQA
HAstV3 LQVLVNSSRKADVYAIQQYLNNQTNNKVQLTSIFLVKVTTSFQVNNYLSYFYRSYGT-GTTVENLKVRSDTTAQDVNFP-
HAstV4 LGVYVGIHKKALVYFMQQYVNPHTNNKHQVSSIFLIKPTENFSITNYMSYFFRESQS-DQNVANLKIRPQTWQQTVNFQR
HAstV5 LLVMKGINKKADVYAAQQYKNSISNNKHQITSVYLIRVKENFQVTNHLSYFFREKVD-TTAVELMKIRPQTYHTTVNFVQ
HAstV6 LMVLLDNHKKADVYAAQQYKNPITNDKQQVTSVFLVRVNEGFQVTNHLSYFYRNSVN-TDAVENIKIRSATRHTTVRFNQ
HAstV7 LQVLVDSRKKADVYAVQQYLNHPSNTKDQLTSIFLVKVTTSFQVNNHLSYFYRAAGT-GTAVENFKIRGATSEQNISFS-
HAstV8 LYARVNNTKKADVYGVQQFKSSHTNNRHQITSVFLVRVTTSFQVINYTSYFIRGAES-GSNVSNLKIRDQTYHTPLQFTQ
FcRn
3E8
PL-2
2D9
4B6
3B4
3H4
β12 Loop 4 β14
597 607 617 627 637 647
HAstV1 GGWYLLTNTSIVDGAMPPGWVWNNVELKTNTAYHMDKGLVHLIMPLPESTQMCYEMLTSIP 217
HAstV2 GEWYLLTSTALKENNLPDAWIWDRVELRSDTPYYADQALAYFITPPPVDSQILFEGSTILP 221
HAstV3 VGWYLMTNTAIFNAPAPPGWIWQNVELLNDTAYLIDRGMMHLIMPPPANTQLLFEMRTSVT 216
HAstV4 GKWYLVTNTAIRNGSPPSGWIWDNMELTNDSIYYADQVLAHFINPPPQNSKLYFEVHTTMP 217
HAstV5 NNWYLLTSTVLHTGSIPSGWVWMNQELMNNENYIVDQGMKHLMTTPPVSSQLYFEMLTSLP 215
HAstV6 GSWYLLTSTVLHTGPPVSGWLWMNQELQNDQAYIIDQGIMHLITPPPVSSQIYFEMATLVP 215
HAstV7 EGWYLMTNTATFNPPAPPGWIWKNVELDNNTPYIVDQGMMHLIMSPPVGTQLLFEMKTTVS 216
HAstV8 GKWYLLTSTVMHDGPTSSGWVWMNQELTNNIAYRVDPGMMYLITPPPAASQLYFELHTVLP 217
FcRn
3E8
PL-2
2D9
4B6
3B4
3H4

**Figure S1**

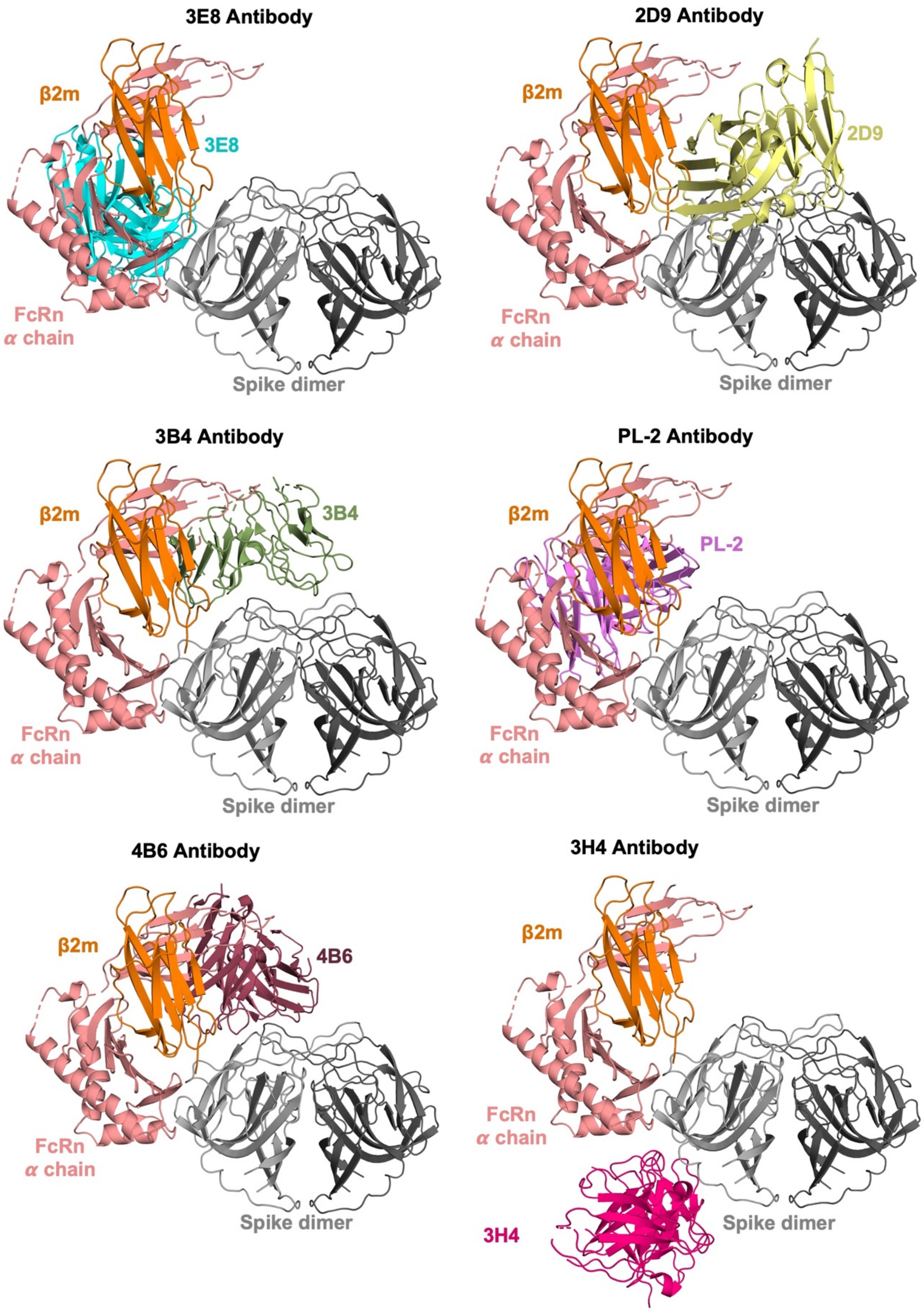
3E8 Antibody
β2m
3E8
FcRn
α chain
Spike dimer
2D9 Antibody
β2m
2D9
FcRn
α chain
Spike dimer
3B4 Antibody
β2m
3B4
FcRn
α chain
Spike dimer
PL-2 Antibody
β2m
PL-2
FcRn
α chain
Spike dimer
4B6 Antibody
β2m
4B6
FcRn
α chain
Spike dimer
3H4 Antibody
β2m
FcRn
α chain
3H4
Spike dimer

**Figure S2**

**Figure Legends:**

**Figure S1. Sequence alignment and epitope mapping of classical HAstV spikes.**

Sequences of spike domain for 8 classical HAstV were aligned using multiple sequence alignment with Clustal Omega. Secondary structure elements (β-sheets) and loop numbering are indicated above the sequences. Black triangles denote spike interface residues mediating FcRn binding; color-coded circles mark neutralizing antibody epitopes, with distinct colors corresponding to specific antibodies.

**Figure S2. Steric competition between neutralizing antibodies and FcRn to the HAstV spike.**

Structural alignment of antibody-spike complexes (3E8 [PDB: 7RK1], 2D9 [PDB: 7RK2], PL-2 [PDB: 5KOV], 3B4 [PDB: 9CBN], 4B6 [PDB: 9CN2], 3H4 [PDB: 9CBN]) with the FcRn-spike complex. Steric clashes between FcRn and neutralizing antibodies (3E8, 2D9, PL-2, 3B4, 4B6) are shown. 3H4 shows no predicted steric interference with FcRn.

**Table S1. Binding affinity parameters for HAstV spike with FcRn.**

| | | Kd (M) | $k_a$ (1/Ms) | $k_{dis}$ (1/s) | $R^2$ |
|---|---|---|---|---|---|
| **pH 7.5** | **HAstV1** | $2.47 \times 10^{-7}$ | $8.75 \times 10^{4}$ | $2.16 \times 10^{-2}$ | 0.994 |
| | **HAstV2** | $8.98 \times 10^{-7}$ | $3.30 \times 10^{4}$ | $2.96 \times 10^{-2}$ | 0.985 |
| | **HAstV6** | $4.70 \times 10^{-7}$ | $3.35 \times 10^{4}$ | $1.57 \times 10^{-2}$ | 0.999 |
| | **HAstV8** | $4.58 \times 10^{-7}$ | $3.22 \times 10^{4}$ | $1.47 \times 10^{-2}$ | 0.995 |
| **pH 6.0** | **HAstV1** | $5.06 \times 10^{-8}$ | $1.25 \times 10^{5}$ | $6.33 \times 10^{-3}$ | 0.979 |
| | **HAstV2** | $4.17 \times 10^{-8}$ | $1.12 \times 10^{5}$ | $4.68 \times 10^{-3}$ | 0.985 |
| | **HAstV6** | $5.20 \times 10^{-8}$ | $7.08 \times 10^{4}$ | $3.69 \times 10^{-3}$ | 0.992 |
| | **HAstV8** | $5.98 \times 10^{-8}$ | $7.21 \times 10^{4}$ | $4.31 \times 10^{-3}$ | 0.989 |

**Table S2. X-ray data collection and refinement statistics.**

| | **FcRn-HAstV2** | **FcRn-HAstV6** |
|---|---|---|
| **Data Collection** | | |
| Beamline | NSLS-II 17-ID-2 | NSLS-II 17-ID-2 |
| Wavelength (Å) | 0.980 | 0.980 |
| Resolution (Å) | 29.52 - 3.07 (3.15 – 3.07) | 29.14 - 2.97 (3.05 – 2.97) |
| Space group | C 1 2 1 | C 1 2 1 |
| Unit cell a, b, c (Å) | 174.9 157.1 86.2 | 211.9 76.7 53.8 |
| α, β, γ (°) | 90 122.1 90 | 90 98.5 90 |
| Total reflections | 262,951 | 124,548 |
| Unique reflections | 37,066 | 17,653 |
| Multiplicity | 7.1 (7.4) | 7.1 (6.9) |
| Completeness (%) | 99.6 (98.7) | 99.6 (96.7) |
| Mean I/sigma(I) | 11.5 (2.1) | 10.7 (2.1) |
| Rsym | 0.14 (0.84) | 0.15 (0.82) |
| Rpim | 0.09 (0.50) | 0.09 (0.51) |
| $CC_{1/2}$ | 0.99 (0.63) | 0.99 (0.73) |
| **Refinement** | | |
| Resolution (Å) | 29.52 - 3.07 | 29.14 - 2.97 |
| Reflections total / Rfree | 37,044 / 2599 | 17,635 / 1292 |
| Rcryst / Rfree | 0.19 / 0.24 | 0.23 / 0.28 |
| No. of copies in ASU | 1 | 1 |
| Number of atoms | 9228 | 4351 |
| macromolecules | 9228 | 4351 |
| solvent | - | - |
| Average B-value($Å^2$) | 75 | 66 |
| Macromolecules | 75 | 66 |
| Spike | 56 | 52 |
| FcRn | 86 | 75 |
| Wilson B ($Å^2$) | 70 | 63 |
| Bond angle (°) | 0.48 | 0.54 |
| Bond length (Å) | 0.002 | 0.002 |
| Favored | 97.6 | 96.8 |
| Allowed | 2.4 | 3.2 |
| **PDB Code** | **9OC7** | **9OC6** |
